# Metabolite profiling reveals organ-specific flavone accumulation in *Scutellaria* and identifies a scutellarin isomer isoscutellarein 8-*O*-β-glucuronopyranoside

**DOI:** 10.1101/2021.09.25.461812

**Authors:** Bryce C. Askey, Dake Liu, Garret M. Rubin, Andrew R. Kunik, Yeong Hun Song, Yousong Ding, Jeongim Kim

**Author notes:** **Corresponding Authors:** Jeongim Kim;, Yousong Ding.

## Abstract

*Scutellaria* is a genus of plants containing multiple species with well-documented medicinal effects. *S. baicalensis* and *S. barbata* are among the best-studied *Scutellaria* species, and previous works have established flavones to be the primary source of their bioactivity. Recent genomic and biochemical studies with *S. baicalensis* and *S. barbata* have advanced our understanding of flavone biosynthesis in *Scutellaria*. However, as over several hundreds of *Scutellaria* species occur throughout the world, flavone biosynthesis in most species remains poorly understood. In this study, we analyzed organ-specific flavone profiles of seven *Scutellaria* species, including *S. baicalensis, S. barbata* and two species native to the Americas (*S. wrightii* to Texas and *S. racemosa* to Central and South America). We found that the roots of almost all these species produce only 4’-deoxyflavones, while 4’-hydroxyflavones are accumulated exclusively in their aerial parts. On the other hand, *S. racemosa* and *S. wrightii* also accumulated high levels of 4’-deoxyflavones in their aerial parts, different with the flavone profiles of *S. baicalensis* and *S. barbata*. Furthermore, our metabolomics and NMR study identified the accumulation of isoscutellarein 8-*O*-β-glucuronopyranoside, a rare 4’-hydroxyflavone, in the stems and leaves of several *Scutellaria* species including *S. baicalensis* and *S. barbata*, but not in *S. racemosa* and *S. wrightii*. Distinctive organ-specific metabolite profiles among *Scutellaria* species indicate the selectivity and diverse physiological roles of flavones.

## Introduction

Medicinal plants have been used in the traditional medicines of indigenous populations for thousands of years. Due to this widespread usage, modern research techniques are being applied to identify the compounds responsible for these medicinal properties and to characterize their modes of action (Shang et al., 2010). A negative consequence of increased attention to and demand for medicinal plants is the endangerment of native plant populations resulting from overharvesting (Cole et al., 2007). Therefore, development of biotechnology-based mass production systems for these medicinal compounds is desirable. Development of effective biotechnology for chemical production requires an understanding of the biosynthesis of the compounds of interest. In this work, we analyze the levels of flavones in various organs of multiple species from the *Scutellaria* genus to better understand flavone biosynthesis in *Scutellaria*.

Part of the mint family Lamiaceae, *Scutellaria* is a genus of plants containing several hundred species with well-documented medicinal effects. Extracts from the aerial parts of *S. barbata* are commonly applied in Eastern medicines to treat swelling, inflammation, and cancer (Tao and Balunas, 2016). These activities, and especially its anticancer effects, have drawn significant attention to *S. barbata*, and early phase clinical trials of aqueous extracts have demonstrated its selective cytotoxicity towards breast cancer cells (Chen et al., 2012). In addition, *S. barbata* extracts have exhibited remarkable activity towards multi-drug resistant strains of bacteria (Tsai et al., 2018). *S. baicalensis* is another species extensively applied in Eastern medicines, with extracts of its roots being prescribed to treat diarrhea, dysentery, hypertension, inflammation, and a variety of other diseases (Zhao et al., 2019b). Numerous clinical studies have demonstrated the neuroprotective, antibacterial, antitumor, antioxidant, and other beneficial health effects of these extracts (Zhu et al., 2016; Saralamma et al., 2017; Tao et al., 2018).

One class of bioactive compounds in *Scutellaria* is flavones (Karimov and Botirov, 2017; Zhao et al., 2019b). *Scutellaria* species produce two classes of flavones: 4’-hydroxyflavones and 4’-deoxyflavones (Fig. 1, Fig. S1). 4’-Hydroxyflavones, including apigenin and its derivatives, are relatively common across the plant kingdom whereas 4’-deoxyflavones, which include chrysin and its derivatives, are relatively rare outside of *Scutellaria* with the exception of several plant species not in the genus (Kato et al., 1992; Rao et al., 2002; Rao et al., 2009). Recent works in *S. baicalensis* and *S. barbata* have identified multiple enzymes responsible for flavone biosynthesis in *Scutellaria*, and have described the differential activity of specific enzymes towards either 4’-hydroxyflavones or 4’-deoxyflavones (Zhao et al., 2016; Zhao et al., 2018; Zhao et al., 2019a). The enzyme selectivity leads to an organ-specific pattern of flavone accumulation. In this pattern, 4’-hydroxyflavones accumulate at higher concentrations in the aerial parts of the plant than in the roots, and the roots contain higher concentrations of 4’-deoxyflavones as compared to the aerial parts (Tao and Balunas, 2016; Xu et al., 2020).

**Figure 1.**
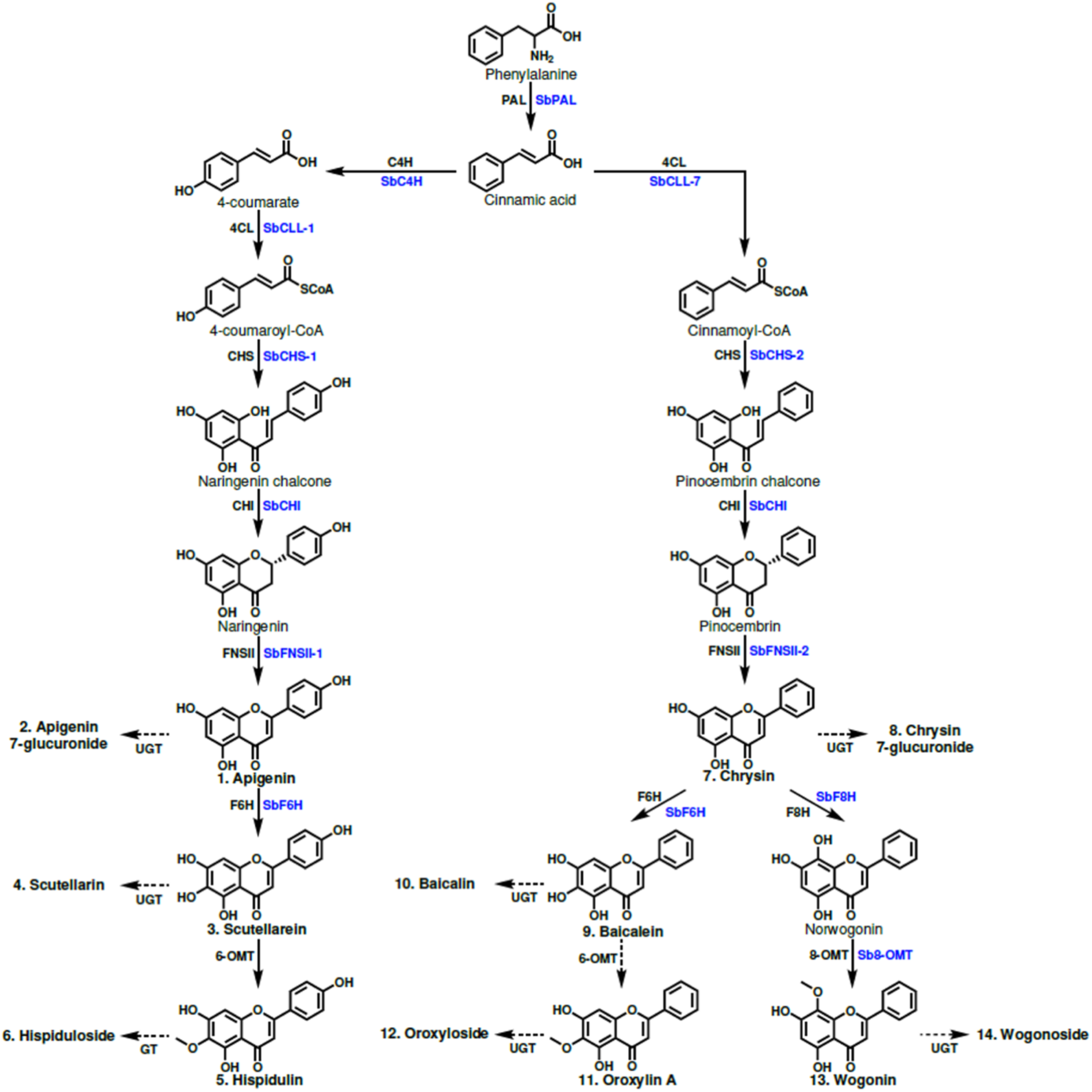
Proposed 4’-hydroxyflavone and 4’-deoxyflavone pathway. Structures of glycosylated flavones are not shown to save space but are included in Fig. S1. Enzyme names in blue are specific isoforms that have been identified in *S. baicalensis*, and enzyme names in black are general names. Flavones that were quantified have names in bold and are numbered to match the labeling of Figure 2. Enzymes are phenylalanine ammonia lyase (SbPAL), cinnamate 4-hydroxylase (SbC4H), cinnamate-CoA ligase (SbCLL-7), 4-coumarate CoA ligase (SbCLL-1), chalcone synthase (SbCHS-1), pinocembrin-chalcone synthase (SbCHS-2), chalcone isomerase (SbCHI), flavone synthase I (SbFNSI), flavone synthase II (SbFNSII), flavone 6-hydroxylase (SBF6H), flavone 8-hydroxylase (SbF8H), and 8-O-methyl transferase (Sb8-OMT).

Flavone profiles of *S. baicalensis* and *S. barbata* have been described and their reference genomes have been established to further support the biosynthetic studies of flavones. However, due to the large number of uncharacterized species in the genus, it is unknown if the overall flavone pathway and the organ-specific accumulation patterns of *S. baicalensis* and *S. barbata* are well-conserved across the genus. In this work, we aimed to expand the current knowledge of flavone diversity in *Scutellaria* by analyzing metabolite profiles of seven species. These species included two well-studied species, *S. baicalensis* and *S. barbata*, and two species native to warm climates, *S. racemosa* and *S. wrightii*. Furthermore, we selected three other *Scutellaria* species widely distributed in Europe, Asia, and North America, including *S. altissima S. tournefortii*, and *S. leonardii* (Hasaninejad et al., 2009; Shang et al., 2010; Sutter et al., 2011), respectively. During this analysis, we unexpectedly identified a 4’-hydroxyflavone which has not been included in recent biosynthetic studies of *S. baicalensis*. We elucidated the structure of this 4’-hydroxyflavone and quantified its level in the seven species. Our results revealed diversity in site and type of flavone accumulated across the species we selected.

## Materials and Methods

### Plant growth conditions

Plants of seven *Scutellaria* species were grown from seed at the University of Florida (Gainesville, Florida, USA) in indoor, climate-controlled conditions at 21-23 °C. Fluorescent lighting of intensity 140 µE m^-2^ s^-1^ was applied in a 16 hour light / 8 hour dark cycle. Plants were watered every 5-8 days, and root, stem, and leaf tissue samples collected in biological triplicate 6-8 weeks after germination. Seeds of all species except for those of *S. racemosa* and *S. wrightii* were obtained from retailers (*S. altissima, S. baicalensis*, and *S. tournefortii*, from Plant World Seeds, and *S. barbata* and *S. leonardii* from Plairie Moon Nursery). To collect seeds of *S. racemosa*, mature plants were taken from a field in Hattiesburg, Mississippi, USA, and grown in indoor, climate-controlled conditions at the University of Florida until seeds were ready to harvest. Seeds of *S. wrightii* were collected directly from mature plants grown in outdoor greenhouse conditions at Far South Wholesale Nursery (Austin, Texas, USA). Herbarium vouchers of all species were submitted to the University of Florida Herbarium, and voucher numbers are provided in Table S3.

### Flavone extraction and quantification

With High Performance Liquid Chromatography (HPLC), 15 flavones were quantified from root, stem, and leaf tissue samples of plants. The flavones quantified included seven 4’-hydroxyflavones, which were apigenin, apigenin-7-glucuronide (apigenin 7-G), scutellarein, scutellarin, hispidulin, hispiduloside, and isoscutellarein-8-glucuronide (isoscutellarein 8-G). The remaining eight flavones were 4’-deoxyflavones, which were chrysin, chrysin-7-glucuronide (chrysin 7-G), baicalein, baicalin, oroxylin A, oroxyloside, wogonin, and wogonoside. The fresh weight of each tissue sample was determined with an analytical balance immediately after harvesting. An extraction solution of 50% HPLC grade methanol was added to each so that the following ratio was achieved: 30 mg tissue/1 mL solvent. Samples were then sonicated for 1 hour at room temperature. Following sonication, the extraction solution was withdrawn and further diluted with additional 50% methanol to achieve a final ratio of 1 mg tissue/1 mL solvent. To remove any remaining particulate, extractions were centrifuged at 15,000 rpm for 5 minutes, and syringe filtered with a filter having a pore size of 0.45 µm.

Flavones were quantified in this final extraction with a Thermo Scientific (Massachusetts, USA) UltiMate 3000 HPLC system. Flavones were separated with a 3 × 100 mm Acclaim RSLC 120 C18 column, and eluted by a mixture of 0.1% formic acid (A) and 100 % acetonitrile (B) with the following gradient: -8 to 0 min, 5% B; 2 min, 25% B; 2 to 6 min, 25% B; 9 min, 50% B; 9 to 11 min, 50% B; 15 min, 95% B; and 15 to 23 min, 95% B. A flowrate of 0.5 mL/min was used and the column oven temperature set to 40°C. Peak areas were measured at wavelength 276 µm. For all flavones except for isoscutellarein 8-G, calibration mixes of 0.1, 0.5, 1, 5, 10, 25, 50, and 100 ppm were used to convert peak areas to concentrations in ppm. Chemical standards used to prepare calibration mixes were purchased from ChemFaces (Wuhan, China) or MilliporeSigma (Massachusetts, USA), and dissolved in dimethylsulfoxide to generate stocks of 1000, 2000, or 4000 ppm. These stocks were then diluted with 50% methanol and mixed to generate calibrations mixes of the varying concentrations. With the peak areas of these calibration mixes and the molecular weight of each metabolite, flavone concentrations in µmol/g fresh weight were calculated. For relative concentration of isoscutellarein 8-G, only peak areas are reported.

### LC-HRMS

LC-HRMS and HRMS/MS experiments were conducted on Thermo Scientific™ Q Exactive Focus mass spectrometer with Dionex™ Ultimate™ RSLC 3000 uHPLC system, equipped with H-ESI II probe on Ion Max API Source. Acetonitrile (B)/Water (A) containing 0.1% formic acid were used as mobile phases. A typical LC program with a 0.5mL/min flow rate included: 10% B for 2 min, 10-95% B in 8.5 mins, 95% B for 2.5 mins, 95 to 10 % B in 0.5 mins, and re-equilibration in 2% B for 2 mins. The eluents from the first 2 mins and last 3 mins were diverted to a waste bottle by a diverting valve. MS1 signals were acquired under the Full MS positive ion mode covering a mass range of m/z 150-2000, with a resolution at 35,000 and a AGC target at 1e6. Fragmentation was obtained using MS2 discovery and Parallel Reaction Monitoring (PRM) mode using an inclusion list of calculated parental ions. Precursor ions were selected in the orbitrap typically with an isolation width of 3.0 m/z and fragmented in the HCD cell with step-wise collision energies (CE) of 20, 25, and 30. For some ions, the isolation width was 2.0 m/z and step-wise CE of 15, 20, and 25 were used.

### NMR analysis

For the NMR analysis, 1.6 mg of compound were dissolved in 40 µl DMSO-d_6_. 1D and 2D spectra were recorded in a 1.7 mm TCI CryoProbe on a Bruker Avance Neo-600 Console system (Magnex 14.1 T/54 mm AS Magnet) at Advanced Magnetic Resonance Imaging and Spectroscopy facility, McKnight Brain Institute, University of Florida. Spectroscopy data were collected and processed using Topspin 4.1.3 software.

### Chemical shifts

^**1**^**H NMR** (600 MHz, DMSO-d_6_): δ_H_ 12.82 (OH, br, s, 5), 10.34 (OH, br, s, 4’), 8.07 (2H, d, J = 8.66 Hz, 2’, 6’), 6.93 (2H, d, J = 8.66 Hz, 3’, 5’), 6.83 (1H, s, 3), 6.29 (1H, s, 6), 5.44 (OH, br, s, 7), 4.82 (1H, J = 7.91 Hz, 1’’), 3.82 (1H, d, J = 9.65 Hz, 5’’), 3.51 (1H, t, J = 9.26 Hz, 4’’), 3.49 (1H, t, J = 8.55 Hz, 2’’), 3.35 (1H, t, J = 9.03 Hz, 3’’). ^**13**^**C NMR** (151 MHz, DMSO-d_6_): δ_c_ 181.72 (C-4), 169.96 (C-6’’), 163.86 (C-2), 161.12 (C-4’), 157.23 (C-7), 156.90 (C-5), 149.19 (C-9), 128.85 (C-2’, C-6’), 125.11 (C-8), 120.96 (C-1’), 115.96 (C-3’, C-5’), 106.25 (C-1’’), 103.35 (C-10), 102.33 (C-3), 98.86 (C-6), 76.00 (C-5’’), 75.20 (C-3’’), 73.69 (C-2’’), 71.41 (C-4’’).

## Results

### Organ-specific flavone diversity across seven *Scutellaria* species

We selected seven species of *Scutellaria* for organ-specific flavone profiling with High Performance Liquid Chromatography (HPLC). These species included *S. altissima, S. baicalensis, S. barbata, S. leonardii, S. racemosa, S. tournefortii*, and *S. wrightii. S. baicalensis* and *S. barbata* have been used in East Asian medicines for thousand years. *S. racemosa* is native to South and Central America (Krings and Neal, 2001), and *S. wrightii* occurs in southwestern regions of North America, such as Texas (Nelson and Goetze, 2010). *S. altissima, S. tournefortii*, and *S. leonardii* are widely distributed in Europe, Asia, and North America (Hasaninejad et al., 2009; Shang et al., 2010; Sutter et al., 2011), but their flavone profiles have not been studied extensively. We grew plants of each species from seed in climate-controlled conditions, and harvested tissue samples from the roots, stems, and leaves of mature plants in biological triplicate. We then quantified concentrations of six 4’-hydroxyflavones (**1**; apigenin, **2**; apigenin 7-glucuronide (apigenin 7-G), **3**; scutellarein, **4**; scutellarin, **5**; hispidulin, **6**; hispiduloside) and eight 4’-deoxyflavones (**7**; chrysin, **8**; chrysin 7-glucuronide (chrysin 7-G), **9**; baicalein, **10**; baicalin, **11**;oroxylin A, **12**; oroxyloside, **13**; wogonin, **14**; wogonoside) in these samples (Fig. 2, Table 1).

**Table 1.** Organ-specific flavone concentrations collected from 7 *Scutellaria* species via High Performance Liquid Chromatography (HPLC). Units for all flavones are µmol / g fresh weight. Data is presented as mean ± standard error, as calculated from samples taken in biological triplicate.

| Species | Organ | Apigenin | Apigenin 7-G | Scutellarein | Scutellarin | Hispidulin | Hispiduloside |
| --- | --- | --- | --- | --- | --- | --- | --- |
| *S. altissima* | Leaves | 0.00 ± 0.00 | 0.28 ± 0.15 | 0.65 ± 0.28 | 4.17 ± 1.22 | 0.00 ± 0.00 | 0.00 ± 0.00 |
| *S. altissima* | Stems | 0.00 ± 0.00 | 0.00 ± 0.00 | 0.00 ± 0.00 | 1.11 ± 0.13 | 0.00 ± 0.00 | 0.00 ± 0.00 |
| *S. altissima* | Roots | 0.00 ± 0.00 | 0.00 ± 0.00 | 0.00 ± 0.00 | 0.00 ± 0.00 | 0.00 ± 0.00 | 0.00 ± 0.00 |
| *S. baicalensis* | Leaves | 0.19 ± 0.03 | 0.13 ± 0.01 | 0.27 ± 0.14 | 0.10 ± 0.10 | 0.00 ± 0.00 | 0.00 ± 0.00 |
| *S. baicalensis* | Stems | 0.12 ± 0.12 | 0.22 ± 0.01 | 1.04 ± 0.43 | 1.76 ± 0.28 | 0.03 ± 0.02 | 0.00 ± 0.00 |
| *S. baicalensis* | Roots | 0.00 ± 0.00 | 0.00 ± 0.00 | 0.00 ± 0.00 | 0.00 ± 0.00 | 0.00 ± 0.00 | 0.00 ± 0.00 |
| *S. barbata* | Leaves | 0.00 ± 0.00 | 0.01 ± 0.01 | 4.59 ± 0.33 | 1.51 ± 0.70 | 0.10 ± 0.02 | 0.00 ± 0.00 |
| *S. barbata* | Stems | 0.00 ± 0.00 | 0.11 ± 0.02 | 1.90 ± 0.23 | 1.13 ± 0.25 | 0.09 ± 0.01 | 0.00 ± 0.00 |
| *S. barbata* | Roots | 0.00 ± 0.00 | 0.00 ± 0.00 | 0.00 ± 0.00 | 0.00 ± 0.00 | 0.00 ± 0.00 | 0.00 ± 0.00 |
| *S. leonardii* | Leaves | 0.11 ± 0.01 | 0.00 ± 0.00 | 0.22 ± 0.04 | 0.00 ± 0.00 | 0.13 ± 0.01 | 0.00 ± 0.00 |
| *S. leonardii* | Stems | 0.02 ± 0.02 | 0.00 ± 0.00 | 0.11 ± 0.01 | 0.16 ± 0.08 | 0.07 ± 0.00 | 0.00 ± 0.00 |
| *S. leonardii* | Roots | 0.06 ± 0.03 | 0.00 ± 0.00 | 0.00 ± 0.00 | 0.82 ± 0.19 | 0.10 ± 0.03 | 0.00 ± 0.00 |
| *S. racemosa* | Leaves | 0.00 ± 0.00 | 0.00 ± 0.00 | 0.00 ± 0.00 | 1.20 ± 0.35 | 0.63 ± 0.18 | 0.00 ± 0.00 |
| *S. racemosa* | Stems | 0.00 ± 0.00 | 0.00 ± 0.00 | 0.00 ± 0.00 | 0.94 ± 0.33 | 0.25 ± 0.06 | 0.06 ± 0.06 |
| *S. racemosa* | Roots | 0.00 ± 0.00 | 0.00 ± 0.00 | 0.00 ± 0.00 | 0.00 ± 0.00 | 0.00 ± 0.00 | 0.00 ± 0.00 |
| *S. tournefortii* | Leaves | 0.00 ± 0.00 | 0.28 ± 0.07 | 0.00 ± 0.00 | 2.98 ± 0.65 | 0.00 ± 0.00 | 0.00 ± 0.00 |
| *S. tournefortii* | Stems | 0.00 ± 0.00 | 0.00 ± 0.00 | 0.00 ± 0.00 | 1.47 ± 0.46 | 0.00 ± 0.00 | 0.00 ± 0.00 |
| *S. tournefortii* | Roots | 0.00 ± 0.00 | 0.00 ± 0.00 | 0.00 ± 0.00 | 0.00 ± 0.00 | 0.00 ± 0.00 | 0.00 ± 0.00 |
| *S. wrightii* | Leaves | 0.00 ± 0.00 | 0.00 ± 0.00 | 0.00 ± 0.00 | 0.13 ± 0.09 | 0.02 ± 0.00 | 0.00 ± 0.00 |
| *S. wrightii* | Stems | 0.00 ± 0.00 | 0.03 ± 0.01 | 0.00 ± 0.00 | 2.10 ± 0.25 | 0.63 ± 0.08 | 0.00 ± 0.00 |
| *S. wrightii* | Roots | 0.00 ± 0.00 | 0.00 ± 0.00 | 0.00 ± 0.00 | 0.00 ± 0.00 | 0.00 ± 0.00 | 0.00 ± 0.00 |

| Species | Organ | Chrysin | Chrysin 7-G | Baicalein | Baicalin | Oroxylin A | Oroxyloside | Wogonin | Wogonoside |
| --- | --- | --- | --- | --- | --- | --- | --- | --- | --- |
| *S. altissima* | Leaves | 0.75 ± 0.07 | 2.62 ± 0.18 | 0.70 ± 0.18 | 0.00 ± 0.00 | 0.00 ± 0.00 | 0.00 ± 0.00 | 0.00 ± 0.00 | 0.00 ± 0.00 |
| *S. altissima* | Stems | 0.04 ± 0.02 | 0.35 ± 0.05 | 0.15 ± 0.05 | 0.47 ± 0.38 | 0.00 ± 0.00 | 0.06 ± 0.06 | 0.34 ± 0.26 | 1.26 ± 0.51 |
| *S. altissima* | Roots | 0.00 ± 0.00 | 0.00 ± 0.00 | 0.07 ± 0.00 | 5.07 ± 0.66 | 0.14 ± 0.02 | 0.64 ± 0.07 | 2.90 ± 0.21 | 2.30 ± 0.26 |
| *S. baicalensis* | Leaves | 4.84 ± 0.53 | 1.45 ± 0.24 | 0.19 ± 0.08 | 0.20 ± 0.11 | 0.00 ± 0.00 | 0.00 ± 0.00 | 0.00 ± 0.00 | 0.00 ± 0.00 |
| *S. baicalensis* | Stems | 0.18 ± 0.04 | 0.09 ± 0.01 | 0.06 ± 0.03 | 0.87 ± 0.87 | 0.00 ± 0.00 | 0.00 ± 0.00 | 0.11 ± 0.11 | 0.02 ± 0.02 |
| *S. baicalensis* | Roots | 0.00 ± 0.00 | 0.30 ± 0.01 | 0.20 ± 0.03 | 32.81 ± 2.22 | 0.23 ± 0.13 | 0.87 ± 0.87 | 3.49 ± 0.18 | 6.43 ± 0.43 |
| *S. barbata* | Leaves | 0.03 ± 0.00 | 0.00 ± 0.00 | 0.00 ± 0.00 | 0.00 ± 0.00 | 0.00 ± 0.00 | 0.00 ± 0.00 | 0.00 ± 0.00 | 0.00 ± 0.00 |
| *S. barbata* | Stems | 0.00 ± 0.00 | 0.00 ± 0.00 | 0.00 ± 0.00 | 0.00 ± 0.00 | 0.00 ± 0.00 | 0.00 ± 0.00 | 0.03 ± 0.02 | 0.00 ± 0.00 |
| *S. barbata* | Roots | 0.00 ± 0.00 | 0.00 ± 0.00 | 1.49 ± 0.04 | 2.59 ± 0.49 | 0.08 ± 0.01 | 0.34 ± 0.18 | 3.88 ± 0.34 | 1.77 ± 0.30 |
| *S. leonardii* | Leaves | 25.34 ± 0.72 | 0.79 ± 0.09 | 0.22 ± 0.03 | 0.07 ± 0.03 | 0.59 ± 0.11 | 0.00 ± 0.00 | 0.07 ± 0.07 | 0.06 ± 0.02 |
| *S. leonardii* | Stems | 5.37 ± 0.41 | 0.91 ± 0.31 | 0.34 ± 0.14 | 0.17 ± 0.08 | 0.37 ± 0.03 | 0.03 ± 0.03 | 1.24 ± 0.09 | 0.10 ± 0.05 |
| *S. leonardii* | Roots | 0.40 ± 0.08 | 0.18 ± 0.04 | 0.12 ± 0.03 | 5.35 ± 0.94 | 1.86 ± 0.31 | 2.18 ± 0.33 | 5.62 ± 1.10 | 0.00 ± 0.00 |
| *S. racemosa* | Leaves | 0.12 ± 0.01 | 0.00 ± 0.00 | 0.34 ± 0.29 | 7.59 ± 1.95 | 14.39 ± 2.58 | 29.39 ± 8.00 | 0.02 ± 0.02 | 0.00 ± 0.00 |
| *S. racemosa* | Stems | 0.03 ± 0.03 | 0.00 ± 0.00 | 0.24 ± 0.22 | 6.36 ± 2.13 | 5.09 ± 1.09 | 15.17 ± 4.14 | 0.44 ± 0.03 | 0.45 ± 0.19 |
| *S. racemosa* | Roots | 0.00 ± 0.00 | 0.00 ± 0.00 | 0.08 ± 0.02 | 6.98 ± 1.20 | 0.44 ± 0.11 | 2.02 ± 0.38 | 1.82 ± 0.06 | 1.60 ± 0.25 |
| *S. tournefortii* | Leaves | 0.49 ± 0.02 | 4.69 ± 0.97 | 0.53 ± 0.13 | 0.00 ± 0.00 | 0.00 ± 0.00 | 0.00 ± 0.00 | 0.00 ± 0.00 | 0.06 ± 0.06 |
| *S. tournefortii* | Stems | 0.01 ± 0.01 | 0.15 ± 0.04 | 0.28 ± 0.07 | 0.08 ± 0.04 | 0.00 ± 0.00 | 0.06 ± 0.03 | 0.13 ± 0.05 | 2.43 ± 0.66 |
| *S. tournefortii* | Roots | 0.00 ± 0.00 | 0.08 ± 0.04 | 0.13 ± 0.06 | 0.93 ± 0.08 | 0.08 ± 0.01 | 1.24 ± 0.14 | 2.33 ± 0.20 | 7.61 ± 0.15 |
| *S. wrightii* | Leaves | 0.82 ± 0.17 | 0.13 ± 0.09 | 0.78 ± 0.18 | 1.21 ± 0.63 | 0.10 ± 0.02 | 0.16 ± 0.05 | 0.00 ± 0.00 | 0.00 ± 0.00 |
| *S. wrightii* | Stems | 0.99 ± 0.37 | 2.23 ± 0.11 | 3.10 ± 1.31 | 29.90 ± 0.92 | 2.24 ± 0.43 | 14.87 ± 0.26 | 0.62 ± 0.07 | 0.40 ± 0.11 |
| *S. wrightii* | Roots | 0.00 ± 0.00 | 0.00 ± 0.00 | 18.10 ± 1.37 | 43.99 ± 9.53 | 1.17 ± 0.38 | 4.22 ± 1.17 | 3.13 ± 0.53 | 3.09 ± 0.87 |

**Figure 2.**
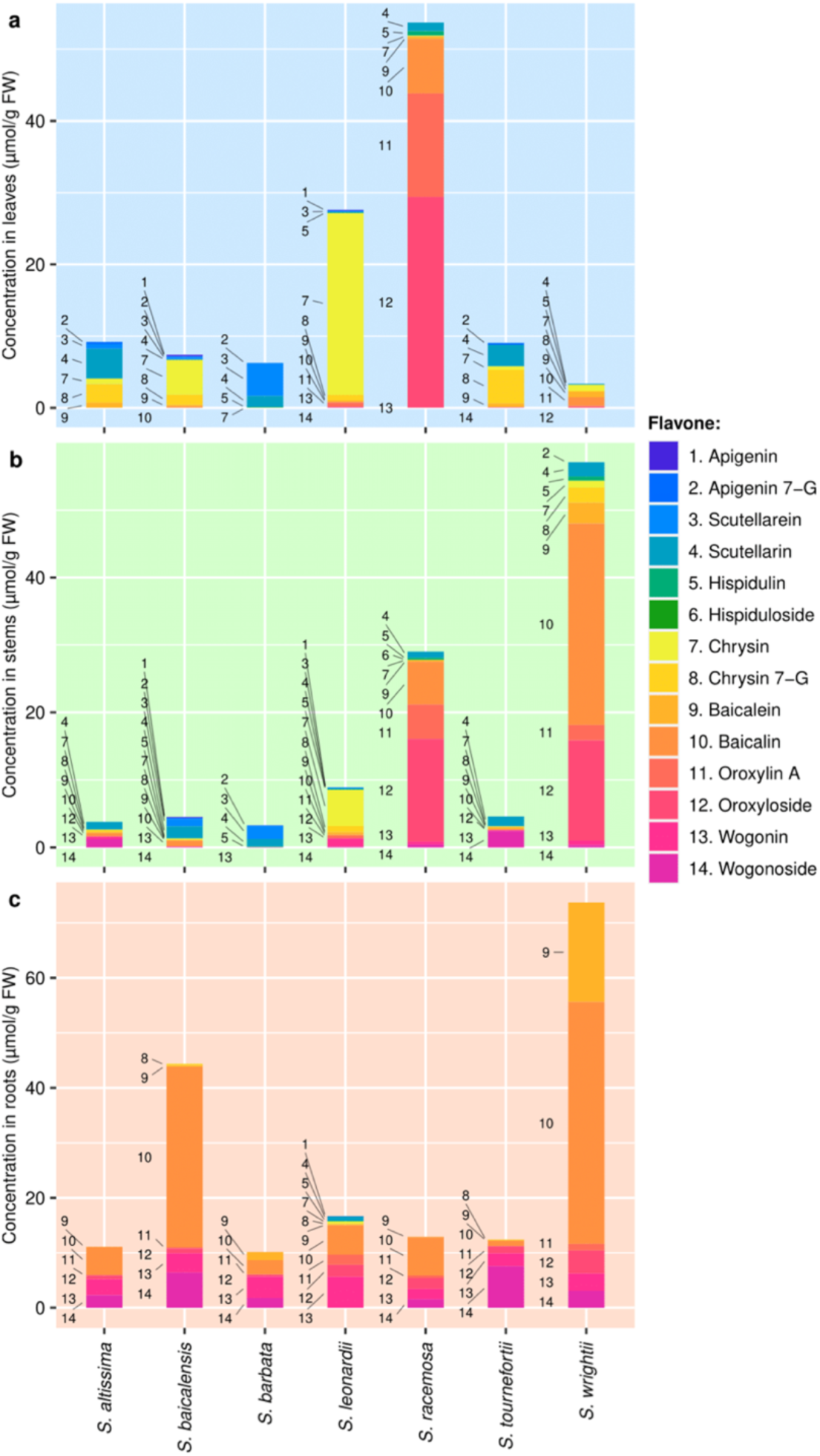
Metabolite data collected from the (a) leaves, (b) stems, and (c) roots of 7 *Scutellaria* species via High Performance Liquid Chromatography (HPLC). Samples were taken in biological triplicate, and the average concentration of each metabolite calculated. Metabolites are numbered to match their order of occurrence in the flavone pathway, shown in Figure 1.

Our root-specific flavone profiling indicated that the 4’-deoxyflavone pathway appears to be very well-conserved across all seven species (Fig. 2c). We detected at least six of eight tested 4’-deoxyflavones in the root of each species (Table 1). Interestingly, although chrysin is proposed to serve as a precursor for all other 4’-deoxyflavones, we found it at a low level (0.40 ± 0.08 µmol / g fresh weight) only in the root of *S. leonardii*, and its glycosylated form, chrysin 7-G, appeared in the roots of three species, *S. baicalensis, S. leonardii*, and *S. tournefortii* (Table 1), ranging from 0.08 to 0.30 µmol / g fresh weight. On the other hand, we observed the accumulation of baicalein, baicalin, oroxylin A, and oroxyloside in the roots of all seven species (Fig. 1). Except for *S. tournefortii*, all species accumulated 1.4 to 12 times more baicalein and baicalin than oroxylin A and oroxyloside (Fig. 2c, Table 1), presumably suggesting the relatively low catalytic activity of 6-OMT (Fig. 1). The highest amount of baicalein (18.10 ± 1.37 µmol / g fresh weight) and baicalin (43.99 ± 9.53 µmol / g fresh weight) was found in the root of *S. wrightii*, followed by *S. baicalensis* (baicalein: 0.20 ± 0.03 µmol / g fresh weight; baicalin 32.81 ± 2.22 µmol / g fresh weight). Interestingly, the root of *S. wrightii* also produced the highest amount of oroxylin A (1.17 ± 0.38 µmol / g fresh weight) and oroxyloside (4.22 ± 1.17 µmol / g fresh weight), while *S. leonardii* was the second best producer. Chrysin can also be converted to wogonin and wogonoside through the reaction of SbF8H and Sb8-OM (Fig. 1). The roots of all species accumulated 1.60 ± 0.25 to 7.61 ± 0.15 µmol / g fresh weight of wogonin and wogonoside, except for *S. leonardii* whose root had no wogonoside. Finally, the absence of 4’-hydroxyflavones in the roots of all but one species (*S. leonardii*) indicates their specificity to the aerial organs of the plant in most species we selected.

Aerial tissue-specific flavone profiles of the selected species were much more varied than root-specific profiles (Fig. 2a, b). First, we detected two to four of six selected 4’-hydroxyflavones in the leaves of all species analyzed, and one to five in their stems (Fig. 2). Of note, this pathway seemed to be inactive in the roots (Figs. 1 and 2). Except for *S. baicalensis* and *S. wrightii*, the leaves of these species accumulated more 4’-hydroxyflavones than the stems. The highest amount of 4’-hydroxyflavones in the leaves was observed in *S. barbata* (6.21 µmol / g fresh weight), followed by *S. altissima* (5.10 µmol / g fresh weight) (Table 1). Among all species, the leaves of *S. wrightii* contained the lowest amount of 4’-hydroxyflavones (0.15 µmol / g fresh weight), while its stems accumulated 2.76 µmol / g fresh weight of these compounds. Among the six selected 4’-hydroxyflavones, we were unable to detect hispidulin, or its glucoside, hispiduloside, in the leaves or stems of two species, *S. altissima* and *S. tournefortii*. Hispiduloside was particularly rare, and out of all tissue samples taken, we only detected it in the stems of *S. racemosa*. Although these more advanced steps in the biosynthetic pathway may not be well-conserved (Fig. 1), the accumulation of scutellarin in the aerial tissues of all seven species indicates at least partial retention of 4’-hydroxyflavone biosynthesis in these species (Fig. 2a, b). Apigenin is a biosynthetic precursor of all other selected 4’-hydroxyflavones. Interestingly, it was scarcely accumulated, as we detected apigenin at low levels (0.02 to 0.19 µmol / g fresh weight) in the aerial tissues of only two species, *S. baicalensis* and *S. leonardii*. This pattern is analogous to the low accumulation of chrysin in our root tissue samples.

In addition to 4’-hydroxyflavones, we observed that several species accumulate one to eight of the selected 4’-deoxyflavones in their aerial parts. Remarkably, the leaves of all seven species accumulated chrysin (0.03 to 25.34 µmol / g fresh weight). Except for *S. barbata* the stems of all sepeceis also produced chrysin (0.01 to 5.37 µmol / g fresh weight). The wide distribution and accumulation of chrysin in the aerial parts are strikingly different with its accumulation in the roots (Figs 2 and 3). Furthermore, *S. racemosa* accumulated 51.85 and 27.78 µmol / g fresh weight of 4’-deoxyflavones in its leaves and stems, higher than their levels in the roots (12.94 µmol / g fresh weight). *S. wrightii* also accumulated a high concentration of 4’-deoxyflavones in their stems (54.35 µmol / g fresh weight), while its roots produced 73.70 µmol / g fresh weight. Of note, *S. leonardii, S. racemosa*, and *S. wrightii* accumulated high concentrations of oroxylin A or oroxyloside in their stems, and *S. racemosa* also in its leaves (Fig. 3, Table 1). This finding is especially remarkable considering the relative rarity of these 4’-deoxyflavones in *S. baicalensis* and *S. barbata*, two well-studied species (Fig. 3). Overall, our detection of chrysin in the leaves of all species analyzed and baicalein in stems and leaves of most species suggests that specificity of 4’-deoxyflavones in roots is less than that of 4’-hydroxyflavones in aerial tissues.

**Figure 3.**
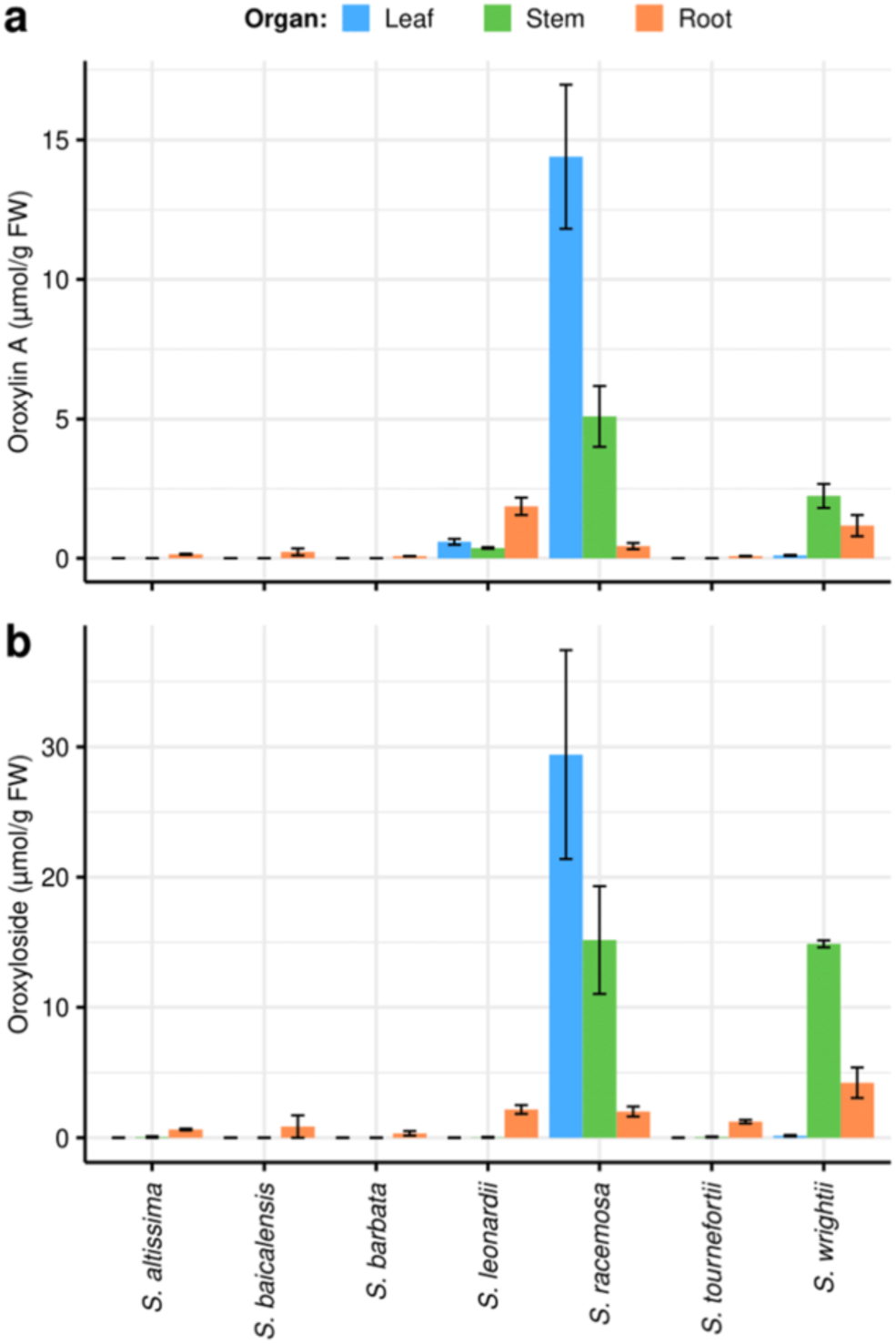
Organ-specific (a) oroxylin A and (b) oroxyloside concentrations in 7 *Scutellaria* species, as determined via High Performance Liquid Chromatography (HPLC). Concentrations were averaged from tissue samples taken from 3 biological replicates, and error bars represent standard error.

### The structural elucidation of a new scutellarin isomer

During our metabolite analysis, we detected multiple new metabolites which we were unable to unambiguously assign their identities. Of these unknown metabolites, one drew our interest because of its pattern of accumulation across the tissue samples we collected (Fig. 4). In our HPLC chromatograms, we detected the peak corresponding to this metabolite in the aerial parts of *S. baicalensis* and *S. barbata*, but not in S. *racemosa*. The peak was absent in root chromatograms collected from all seven species. The aerial specificity of this unknown metabolite led us to hypothesize that it was a 4’-hydroxyflavone. To elucidate its structure, we analyzed the unknown metabolite from our *S. barbata* leaf extracts by the liquid chromatography-high resolution mass spectrometry (LC-HRMS). Interestingly, its molecular weight was identical to scutellarin ([M + H]^+^ *m/z* 463.0866, calculated for C_21_H_19_O_12_^+^, 463.0871), but they were eluted with different retention times (t_m_ = 6.28 min for scutellarin vs 6.94 min for the unknown compound)(Fig. 5a). Furthermore, they gave rise to the same major MS/MS fragment, suggesting them to be two isomers (Fig. S2).

**Figure 4.**
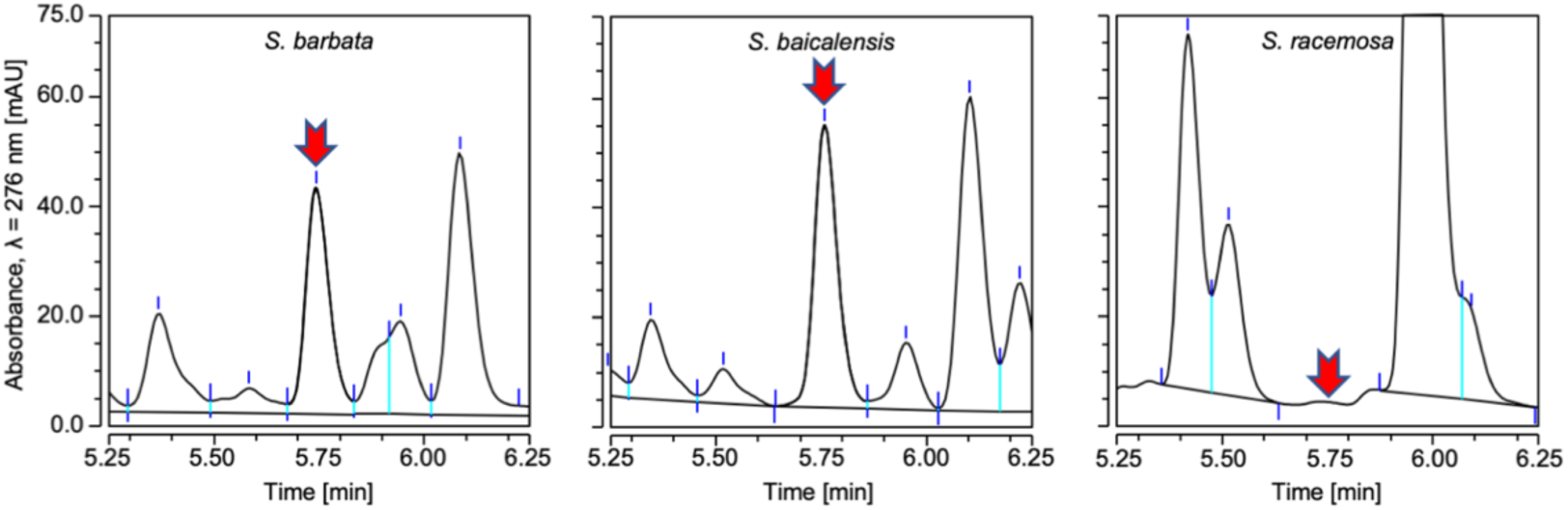
Comparison of chromatograms collected via HPLC from *S. barbata, S. baicalensis*, and *S. racemosa* stems. Time interval displayed was selected to center the unknown peak in the chromatograms.

**Figure 5.**
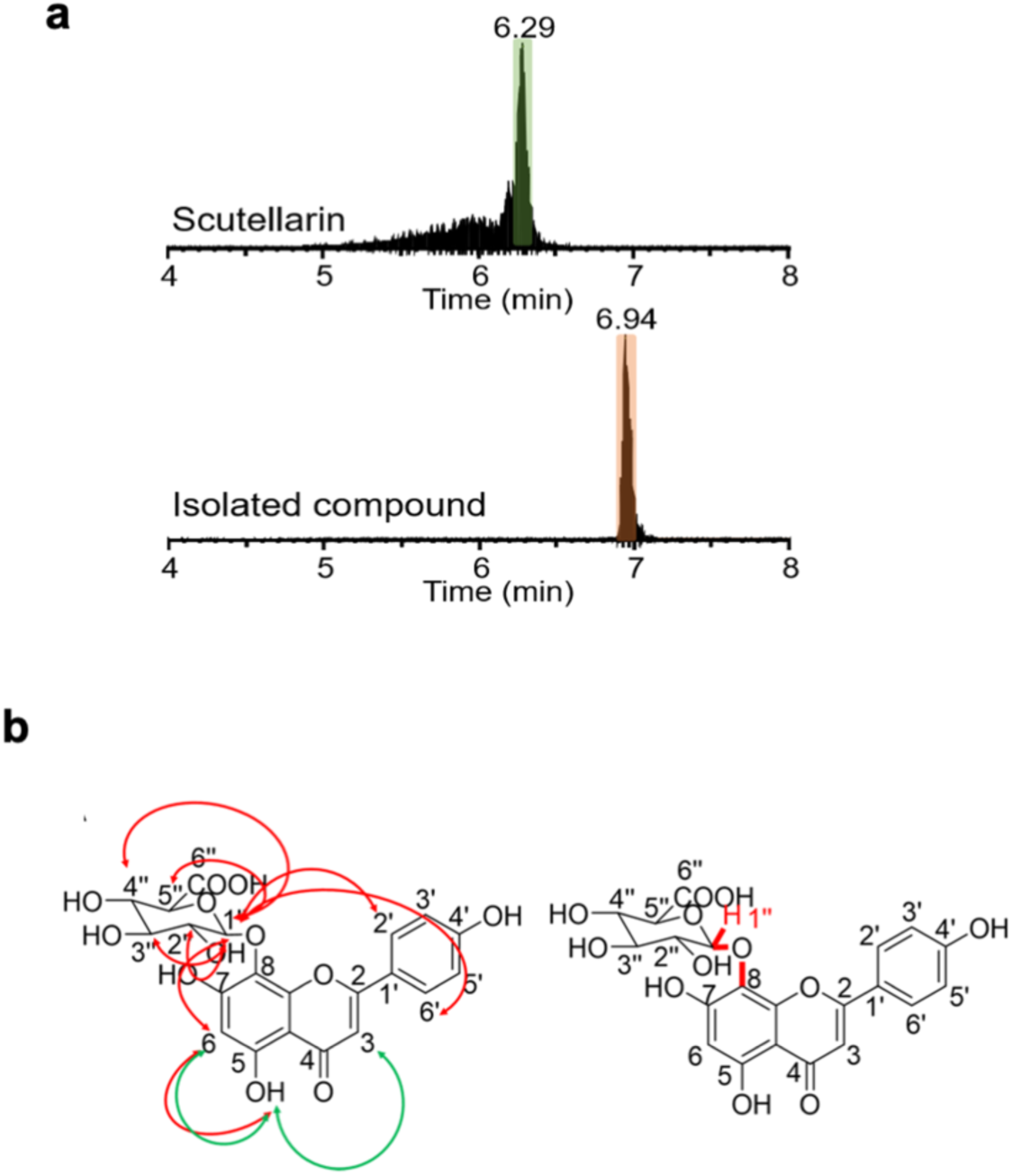
Characterization of a new scutellarin isomer. (a). Standard scutellarin and isolated compound were eluted with different retention times in LC-HRMS analysis. (b). Key NMR correlations of isolated compound. 1D-NOESY and1D-ROESY correlations are represented by red and green two-way arrows, respectively (left). A three-bond HMBC correlation from H-1’’ to C-8 (right). metabolite.**+** NMR data used to elucidate structure of unknown metabolite.

To further elucidate the structure of this compound, we performed 1D and 2D NMR analysis (Figs. S3-5). Comparison of its ^1^H and ^13^C chemical shifts to those of scutellarin allowed the assignment of D-glucuronide (C1’’ to 6’’), 1,4-disubstituted benzene ring (C1’ to 6’), and the carbons on the flavone ring (Jiang et al., 2016)(Fig. 5b, Table S1). Based on the ^1^H chemical shift and coupling constant of the anomer proton H-1’’ (*J* = 7.86 Hz), the glucuronyl moiety was determined to be on the β configuration (Ko et al., 2018). Compared with scutellarin, the aromatic proton at δ_H_ 6.99 (1H, s) was initially assigned to H-8 of the flavone. However, according to 1D-selective nuclear overhauser effect spectroscopies (NOSEY, resonance frequency at 6.28 ppm or 12.81 ppm) and an 1D-selective rotating frame overhauser enhancement spectroscopy (ROSEY, resonance frequency at 12.81 ppm), OH-5 correlates with H-3 (δ = 6.83) and a proton at δ = 6.29 (Fig. 5b), leading to the assignment of this proton at 6 position (δ = 6.29 ppm). This assignment was further supported by the weak NOE effects of H-6 with OH-5 and H-1’’ on the glucuronyl moiety, which further indicated the proximity of the glucuronyl moiety at 7 or 8 position. H-1’’ also showed weak NOE effects with H2’ and H6’ on the 1,4-disubstituted benzene ring, suggesting the potential configuration of the compound, where the glucuronyl moiety could be close to the aromatic system. According to an HMBC correlation from H1’’ to C8 of the flavone, we then definitely assigned the glucuronyl moiety at 8 position (Fig. 5b). Collectively, our 1D and 2D NMR analysis revealed the isolated compound as isoscutellarein 8-*O*-β-glucuronopyranoside (isoscutellarein 8-G). Comparison with the reported ^1^H and ^13^C chemical shifts of this compound (Billeter et al., 1991) confirmed this structural determination (Table S2).

After confirming the identity of this unknown metabolite as isoscutellarein 8-G, we then quantified its relative abundance in all organ-specific tissue samples we collected (Fig. 6, Table 2). Isoscutellarein 8-G was accumulated only in the aerial parts of all species, matching the pattern which we had previously observed for 4’-hydroxyflavones including scutellarin. *S. barbata* accumulated the greatest overall concentrations of isoscutellarein 8-G in its stems. *S. baicalensis, S. altissima*, and *S. tournefortii* also accumulated isoscutellarein 8-G in their stems. In contrast, *S. leonardii, S. racemosa*, and *S. wrightii* accumulated no isoscutellarein 8-G in their aerial parts. It is noteworthy that these three species accumulated oroxylin A and its glycoside in their aerial parts (Fig. 3).

**Table 2.** Organ-specific isoscutellarein 8-G peak areas collected from 7 *Scutellaria* species via High Performance Liquid Chromatography (HPLC). Data is presented as mean ± standard error, as calculated from samples taken in biological triplicate.

| Species | Organ | Isoscutellarein 8-G |
| --- | --- | --- |
| *S. altissima* | Leaves | 0.00 ± 0.00 |
| *S. altissima* | Stems | 0.26 ± 0.01 |
| *S. altissima* | Roots | 0.00 ± 0.00 |
| *S. baicalensis* | Leaves | 0.10 ± 0.05 |
| *S. baicalensis* | Stems | 2.17 ± 0.60 |
| *S. baicalensis* | Roots | 0.00 ± 0.00 |
| *S. barbata* | Leaves | 2.34 ± 0.49 |
| *S. barbata* | Stems | 3.89 ± 0.60 |
| *S. barbata* | Roots | 0.00 ± 0.00 |
| *S. leonardii* | Leaves | 0.00 ± 0.00 |
| *S. leonardii* | Stems | 0.00 ± 0.00 |
| *S. leonardii* | Roots | 0.00 ± 0.00 |
| *S. racemosa* | Leaves | 0.00 ± 0.00 |
| *S. racemosa* | Stems | 0.00 ± 0.00 |
| *S. racemosa* | Roots | 0.00 ± 0.00 |
| *S. tournefortii* | Leaves | 0.18 ± 0.03 |
| *S. tournefortii* | Stems | 0.90 ± 0.24 |
| *S. tournefortii* | Roots | 0.00 ± 0.00 |
| *S. wrightii* | Leaves | 0.00 ± 0.00 |
| *S. wrightii* | Stems | 0.00 ± 0.00 |
| *S. wrightii* | Roots | 0.00 ± 0.00 |

**Figure 6.**
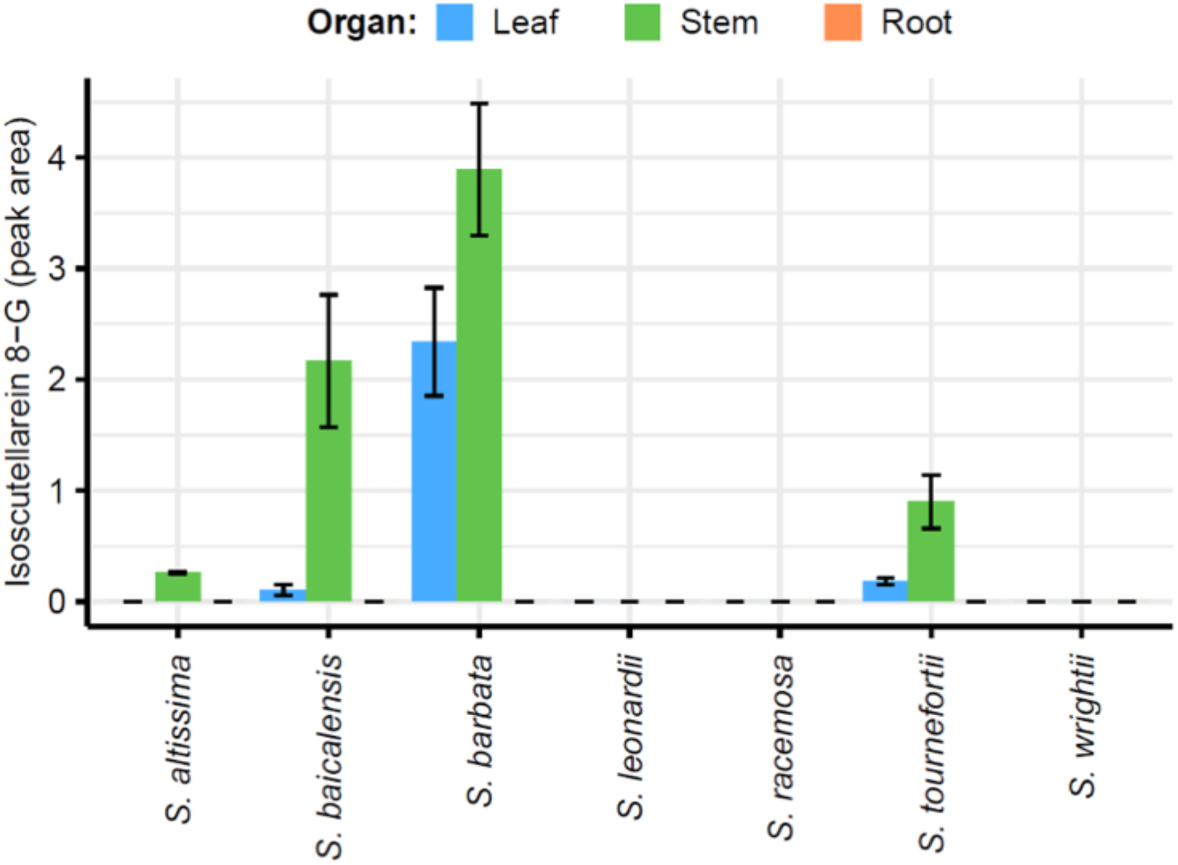
Organ-specific isoscutellarein 8-glucuronide peak areas in 7 *Scutellaria* species, as determined via High Performance Liquid Chromatography (HPLC). Peak areas were averaged from tissue samples taken from 3 biological replicates, and error bars represent standard error.

## Discussion

From our analysis of organ-specific flavone diversity, we determined profiles for *S. baicalensis* and *S. barbata*, which matched closely with previous publications (Zhao et al., 2016; Xu et al., 2020). In these flavone profiles, high concentrations of 4’-deoxyflavones accumulated in the roots, and much lower concentrations of 4’-deoxyflavones and 4’-hydroxyflavones accumulated in the stems and leaves (Fig 2, Table 1). As described by Q. Zhao et al. (2016), the root-favored accumulation of 4’-deoxyflavones by *S. baicalensis* is due to root-specific overexpression of several enzyme isoforms with activity exclusively, or near exclusively in 4’-deoxyflavone biosynthesis (Zhao et al., 2016). In contrast to the pattern we observed in *S. baicalensis* and *S. barbata*, we identified that *S. racemosa* and *S. wrightii* accumulated higher concentrations of 4’-deoxyflavones in their aerial parts as compared to their roots (Fig 2, Table 1). Also, all seven species accumulated chrysin and/or chrysin 7-glucuronide in their leaves (Fig 2a, Table 1). This suggests that the expression of 4’-deoxyflavone enzyme isoforms is not perfectly root-specific, and some enzymes having activities toward 4’-deoxyflavone precursors such as SbCLL-7 and SbCHS-2 may be active in both roots and aerial parts at least under our growth conditions. It is also possible that some fraction of 4’-deoxyflavones are synthesized in the roots and then transported to the aerial parts. The fact that 4’-hydroxyflavones were not detected in roots of most species indicates the selectivity of enzymes towards either 4’-deoxyflavones or 4’-hydroxyflavones (or their respective precursors), as well as organ-specific regulation of biosynthetic gene expression.

We found that *S. racemosa* accumulates the highest concentrations of oroxylin A, and its 7-glucuronide, oroxyloside, in its leaves, among all organs of all species (Fig. 3, Table 1). *S. wrightii* also accumulated notable amounts of oroxylin A and oroxyloside in its stem, but not in its leaves. Oroxylin A is a 4’-deoxyflavone which has been demonstrated to exhibit memory enhancement and neuroprotective effects in rat models (Jeon et al., 2011; Jeon et al., 2012). The most likely route for oroxylin A biosynthesis is the methylation of baicalein at its 6-OH group (Fig. 1) (Elkin et al., 2018). Although previous works have identified a variety of *O*-methyltransferases (OMTs) in plants, the enzymes with high specificity for the 6-OH group in flavonoids are rare, as this reaction is biochemically unfavorable (Zhang et al., 2016a). Work in sweet basil (*Ocimum basilicum*), a species also in the Lamiaceae family with *Scutellaria*, identified a methyltransferase capable of specific methylation of the 6-OH group of scutellarein (Berim et al., 2012). Scutellarein is a 4’-hydroxyflavone identical in structure to baicalein apart from its 4’-OH group. To ensure the proper orientation of its substrate, and thus its regioselectivity, the *O. basilicum* OMT uses a threonine residue to form a hydrogen bond with the 4’-OH group of scutellarein. However, as baicalein has no 4’-OH group, it would be impossible for a regioselective OMT in *S. racemosa* or *S. wrightii* to rely on this interaction during the methylation of baicalein. Research by Zhang et al. (2016) in a liverwort species (*Plagiochasma appendiculatum*) identified a methyltransferase (PaF6OMT) that is capable of methylation of the 6-OH group in baicalein (Zhang et al., 2016b). As this OMT has not yet been structurally characterized, how it achieves its specificity remains unknown. Future work in *S. racemosa* and *S. wrightii* should be directed towards characterizing its biosynthesis of oroxylin A, with specific attention paid to the potential specialization of OMTs in the pathway. Overall, *S. racemosa* and *S. wrightii* are promising targets for biotechnology improvement due to the significant bioactive effects of oroxylin A and oroxyloside. Considering that both species occur in warm area (Texas and South America) (Krings and Neal, 2001; Nelson and Goetze, 2010), accumulation of oroxylin A and oroxyloside in these species may indicate the physiological relevance of oroxylin A and oroxyloside in these species.

Isoscutellarein 8-G was first detected in the liverwort species *Marchantia berteroana* (Markham and Porter, 1975). Following this initial report, Miyaichi et al. detected the flavone in the aerial parts of *S. indica* and *S. baicalensis* (Miyaichi et al., 1988a; Miyaichi et al., 1988b). Aside from these works, few other studies have reported isoscutellarein 8-G in *Scutellaria*, though several have detected its aglycone and 7-*O*-glycosylated forms (Karimov and Botirov, 2017). This rarity in detection may be a result of its low abundance relative to other glycosylated flavones in *Scutellaria*. A potential reason for this low abundance is its unique glycosylation at the 8-*O* position. Flavone 7-*O* glycosylation is more common in *Scutellaria* due to the presence of a hydroxyl group at the 7-*O* position in all flavones synthesized via the core flavone pathway (Fig. 1). On the other hand, 8-*O* glycosylation first requires the activity of an 8-hydroxylase to add the free hydroxyl group to which the carbohydrate will be attached. As the purpose of glycosylation is typically to increase the stability of the flavone for long term storage (Slámová et al., 2018), it’s possible that 8-*O* glycosylation provides slightly greater stability as compared to 7-*O* glycosylation. Therefore, it would be preferable to glycosylate isoscutellarein at the 8-*O* position, even though a free hydroxyl group is also present at the 7-*O* position. Several species may have evolved regioselective glycosyltransferase enzymes for this purpose. Researchers working with a glycosyltransferase from *Bacillus cereus* demonstrated that a single amino acid substitution could alter the primary site of quercetin glycosylation with high specificity (Chiu et al., 2016). Perhaps a similar mutation occurs in several *Scutellaria* species to allow the biosynthesis of isoscutellarein 8-G. Alternatively, it’s possible that the glycosyltransferase enzymes of these species which accumulate isoscutellarein 8-G have less strict regioselectivity, and are capable of glycosylation at both 7-G and 8-G positions. Quantification of isoscutellarein 7-G alongside isoscutellarein 8-G would provide valuable insight regarding these theories. Based on current understanding of flavone biosynthesis, we propose a possible route of isoscutellarein and isoscutellarein 8-G production from apigenin (Fig. S6). Further organ-specific transcriptome study is required to identify enzymes responsible for of isoscutellarein and isoscutellarein 8-G production.

Our quantification of isoscutellarein 8-G across the seven *Scutellaria* species we analyzed revealed an intriguing pattern. Isoscutellarein 8-G was entirely absent in the species of *S. leonardii, S. racemosa*, and *S. wrightii*, all of which accumulate high concentrations of 4’-deoxyflavones such as oroxylin A and oroxyloside in their aerial parts. This specific example is representative of a broader pattern - species with high accumulation of 4’-deoxyflavones in their aerial parts accumulated low concentrations of 4’-hydroxyflavones. This substitution of 4’-hydroxyflavones with 4’-deoxyflavones potentially indicates an evolution to utilize 4’-deoxyflavones to fulfill the physiological roles which 4’-hydroxyflavones do in other species. Works in species outside of *Scutellaria* have demonstrated the anti-herbivory effects of several 4’-hydroxyflavones we quantified here (Sosa et al., 2004; Gallon et al., 2019). However, little is known about the physiological role that 4’-deoxyflavones play in plants. Further research should be devoted to exploring the role of 4’-deoxyflavones in plant growth and stress response to better understand the evolutionary advantage their biosynthesis and accumulation offers.

## Acknowledgments

This work was supported by the United States Department of Agriculture (USDA)-National Institute of Food and Agriculture Hatch project (005681), a startup fund from the Horticultural Sciences Department and Institute of Food and Agricultural Sciences at the University of Florida to J.K, and by NIH (R35 GM128742) to Y.D. NMR studies were performed in the McKnight Brain Institute at the National High Magnetic Field Laboratory’s AMRIS Facility, which is supported by the National Science Foundation Cooperative Agreement No. DMR-1644779, the State of Florida, and an NIH award, S10RR031637. We thank John B. Nelson at A.C. Moore Herbarium and the late William Mark Whitten at the UF for collecting *S. racemosa* in the field. We also thank Dr. Sangtae Kim for the discussion and Dr. Swathi Nadakuduti for scutellarin standard.

## Contributions

B.C.A., Y.D., and J.K. designed the research project; B.C.A., D.L., G.M.R, and Y.S. performed the experiments and analyzed the data; B.C.A., Y.D., and J.K. wrote the manuscript.

## TABLES

Table 1. Organ-specific flavone concentrations collected from 7 *Scutellaria* speces via High Performance Liquid Chromatography (HPLC).

Table 2. Organ-specific isoscutellarein 8-G peak areas collected from 7 *Scutellaria* species.

## Supplemental materials

**Figure S1**. Proposed 4’-hydroxyflavone and 4’-deoxyflavone pathway with structures of glycosylated flavones included. Enzyme names in blue are specific isoforms that have been identified in *S. baicalensis*, and enzyme names in black are general names. Flavones that were quantified have names in bold and are numbered to match the labeling of Figure 2.

**Figure S2**. MS (a), and MS/MS (b) spectra of standard scutellarin and isolated compound.

**Figure S3**. ^1^H NMR spectrum of isolated compound (600 MHz, DMSO-d_6_). Water signals were suppressed by presaturation.

**Figure S4**. ^13^C NMR spectrum of isolated compound (151 MHz, DMSO-d_6_).

**Figure S5**. 2D NMR spectra of isolated compound. **A:**1H-1H COSY; **B**: HSQC; **C:** HMBC; **D**: 2D-NOESY. Positive and negative contours are highlighted in black and green, respectively.

**Figure S6**. Proposed pathway for biosynthesis of isoscutellarein 8-glucuronide in *Scutellaria*.

**Table S1**. Comparison of ^1^H (600 MHz, DMSO-d_6_) chemical shifts of the and a previous literature of scutellarin

**Table S2**. Comparison of ^1^H (600 MHz, DMSO-d_6_) and ^13^C (151 MHz, DMSO-d_6_) chemical shifts of isolated compound and isoscutellarein 8-*O*-β-glucuronopyranoside

**Table S3**. Voucher information for the species used in this study

